# The Ventral Midline Thalamus Mediates Successful Deliberation by Coordinating Prefrontal and Hippocampal Neural Activity

**DOI:** 10.1101/2021.06.30.450519

**Authors:** John J. Stout, Henry L. Hallock, Suhaas S. Adiraju, Amy L. Griffin

## Abstract

When faced with difficult choices, the possible outcomes are considered through a process known as deliberation. In rats, deliberation can be reflected by pause-and-reorienting behaviors, better known as Vicarious Trial and Errors (VTEs). While VTEs are thought to require medial prefrontal cortex (mPFC) and dorsal hippocampal (dHPC) interactions, no work has demonstrated such a dual requirement. The nucleus reuniens (Re) of the ventral midline thalamus is anatomically connected with both the mPFC and dHPC, is required for HPC-dependent spatial memory tasks, and is critical for mPFC-dHPC neural synchronization. Currently, it is unclear if, or how, the Re is involved in deliberation. Therefore, by examining the role of the Re on VTE behaviors, we can better understand the anatomical and physiological mechanisms supporting deliberation. Here, we examined the impact of Re suppression on VTE behaviors and mPFC-dHPC theta synchrony during well-learned performance of a HPC-dependent delayed alternation (DA) task. Pharmacological suppression of the Re increased VTE behaviors that resulted in erroneous choices. These errors were best characterized as perseverative behaviors, where at the decision-point, some rats repeatedly turned in the direction that previously yielded no reward, while simultaneously deliberating. These ‘failed’ deliberations were associated with a reduction in mPFC-dHPC theta coherence at the choice-point. Importantly, the reduction in mPFC-dHPC theta synchrony observed during VTE behaviors was almost entirely driven by Re-suppression induced changes to the mPFC theta oscillation. Our findings suggest that the Re is important for successful deliberation, and mPFC-dHPC interactions, by coordinating local theta oscillations in the mPFC.

## Introduction

When confronted with difficult decisions, the available outcomes are considered through a process known as deliberation. In humans, deliberation can be studied through self-report. However, deliberation in non-human animals is more difficult to study. Nonetheless, research dating back to the 1930s has reported what appear to be deliberative behaviors in rats. When rats are confronted with a ‘difficult’ choice such as a choice that affects whether or how the rat will be rewarded, they sometimes pause, and look ahead towards possible outcomes. These behaviors were defined as *vicarious trial-and-errors* (VTEs) (Muenzinger and Gentry 1931, Muenzinger 1938; Tolman 1939; Tolman 1948; Amsel 1993; Hu & Amsel 1995; Bimonte & Denenberg 2000; Johnson & Redish 2007; Bett et. al., 2012; Gardner et al., 2012; Schmidt et al., 2013; Redish 2016) and have since been identified in humans (Voss et al., 2011; Voss and Cohen, 2017; Santos-Pata and Verschure, 2018).

Since VTEs were first described, it has been demonstrated that VTEs predominately occur during the learning of hippocampal-dependent place-memory tasks (Hu and Amsel, 1995; Bett et al., 2012; Gardner et al., 2012; Schmidt et al., 2013). As rats become familiar with the rules of a place-memory paradigm, VTEs diminish (Papale et al., 2012). However, if rats are required to maintain task-relevant information (e.g. use working-memory), VTEs emerge during decision-making. Manipulating task parameters to induce VTEs has provided critical insight into the brain-circuitry that supports these behaviors (Redish, 2016). For example, inactivation of the hippocampus (HPC) during learning prevents acquisition of place-memory rules, and reduces VTE behaviors (Hu and Amsel, 1995). Moreover, when working-memory demand is applied to place-memory tasks, the medial prefrontal cortex (mPFC) and HPC are both required for successful decision-making (Lee and Kesner, 2003; Wang and Cai, 2006; Wang and Cai, 2008; Churchwell & Kesner 2011;). Importantly, inactivation of the mPFC also impacts VTE behaviors (Schmidt et. al., 2019; Kidder et al., 2020). Thus, both the mPFC and HPC support deliberation in the rat.

Using place-memory paradigms, it was demonstrated that theta-frequency (4-12Hz) fluctuations in the local-field potential (LFP) become synchronized between the mPFC and HPC during decision-making (Jones and Wilson, 2005; Benchenane et. al., 2010; Sigurdsson et al., 2010; O’Neill et al., 2013; Hallock et. al., 2016). Importantly, theta synchrony is strongly linked to *successful* decision-making and increases with task-acquisition (Benchenane et al., 2010; Hyman et al., 2010; Hallock et al., 2016). Interestingly, this relationship is opposite to the relationship between learning and VTE behaviors, with VTE behaviors becoming less frequent as rats reach asymptotic performance on the task (Hu & Amsel 1985; Redish, 2016). Together, these findings suggest that VTE behaviors and theta-synchrony are related, and support the hypothesis that mPFC-HPC communication is related to deliberation (Redish, 2016).

While studies have separately examined the impact of mPFC or HPC inactivation on VTE behaviors (Hu and Amsel 1995; Bett et. al., 2012; Schmidt et. al., 2019; Meyer-Mueller et al., 2020), the anatomical pathways by which the HPC and mPFC are linked to VTEs remains unclear. Although there are multiple routes by which the mPFC and HPC can interact, one of the most prominent projection pathways connecting the dorsal hippocampus (dHPC) with the mPFC is the nucleus reuniens (Re) of the ventral midline thalamus (see Griffin, 2015; Eichenbaum et al., 2017; Dolleman-van der Weel et al., 2019 for reviews). The Re is bi-directionally connected with the mPFC and HPC (Vertes et. al., 2006; Vertes et. al., 2007), and sends a collateral projection to both regions (Varela et al., 2014; Vertes et. al., 2007). Given this anatomical connectivity, the Re is a strong candidate region to coordinate mPFC-HPC communication. In the past decade, it has been demonstrated that the Re is critical for mPFC-HPC dependent tasks (Hembrook and Mair, 2011; Hembrook et al., 2012; Hallock et al., 2013; Layfield et al., 2015; Hallock et al., 2016; Maisson et al., 2018), and supports mPFC-HPC interactions (Ito et al., 2015; Hallock et al., 2016; Ferraris et al., 2018). However, no work to-date has examined *if* the Re contributes to VTEs, and, if so, how this contribution relates to Re’s role in mPFC-dHPC theta synchronization. Therefore, understanding the mechanisms by which the Re contributes to VTEs is likely critical to understanding the neural circuitry that supports deliberation. Our hypothesis is that the Re coordinates mPFC-HPC communication in support of deliberative processes. Under this hypothesis, we predicted that Re suppression would impact the behavioral manifestation of VTEs.

Here, we assessed the impact of Re inactivation on VTEs in a previously published dataset from our lab (Hallock et al., 2016). We found that Re inactivation negatively impacted the *success* of deliberation, or the probability that VTE behaviors led to a correct decision. Moreover, we find that theta coherence, a metric that quantifies the strength of the relationship between two LFP signals in the theta (5-10Hz) range, was reduced under Re suppression during VTE behaviors, but not during non-VTE behaviors. This reduction in theta synchrony was driven by Re contributions to the prefrontal theta oscillation. We then used a separate dataset to show that failed deliberation and reduced theta oscillations in the mPFC under Re inactivation were accompanied by error patterns that are typically observed in the early stages of learning in undisturbed rats (Hallock et al., 2013). We take our findings to indicate that Re inactivation induced brain systems that normally compete with mPFC and HPC during cognition to dominate the decision process.

## Methods

### Subjects

Subjects were 15 adult male Long-Evans hooded rats. Seven subjects whose data was previously published (Hallock et al., 2016) were used for their video-tracking data, mPFC/dHPC recordings, and Re manipulation. Training data from 8 rats whose data was previously published (Hallock et al., 2013) were used in the analyses performed for the dataset shown in **Fig. 4E and Fig. 4F**. Procedures were approved by the University of Delaware Institutional Animal Care and Use Committee.

### Behavioral apparatus and testing room

The behavioral training and testing was recorded on an modified T-maze (Central stem: 116 × 10 cm, goal-arms (2): 56.5 × 10 each, return-arms (2): 112 × 10 cm each). The “start box”, where rats were confined between trials with a wooden barricade, was located at the base of the maze stem (**Fig. 1A**). All training, testing, and recording was performed in a dimly lit room surrounded by black curtains, with visual cues (red and green-tape strips, triangles, and patterned circles).

**Figure 1:**
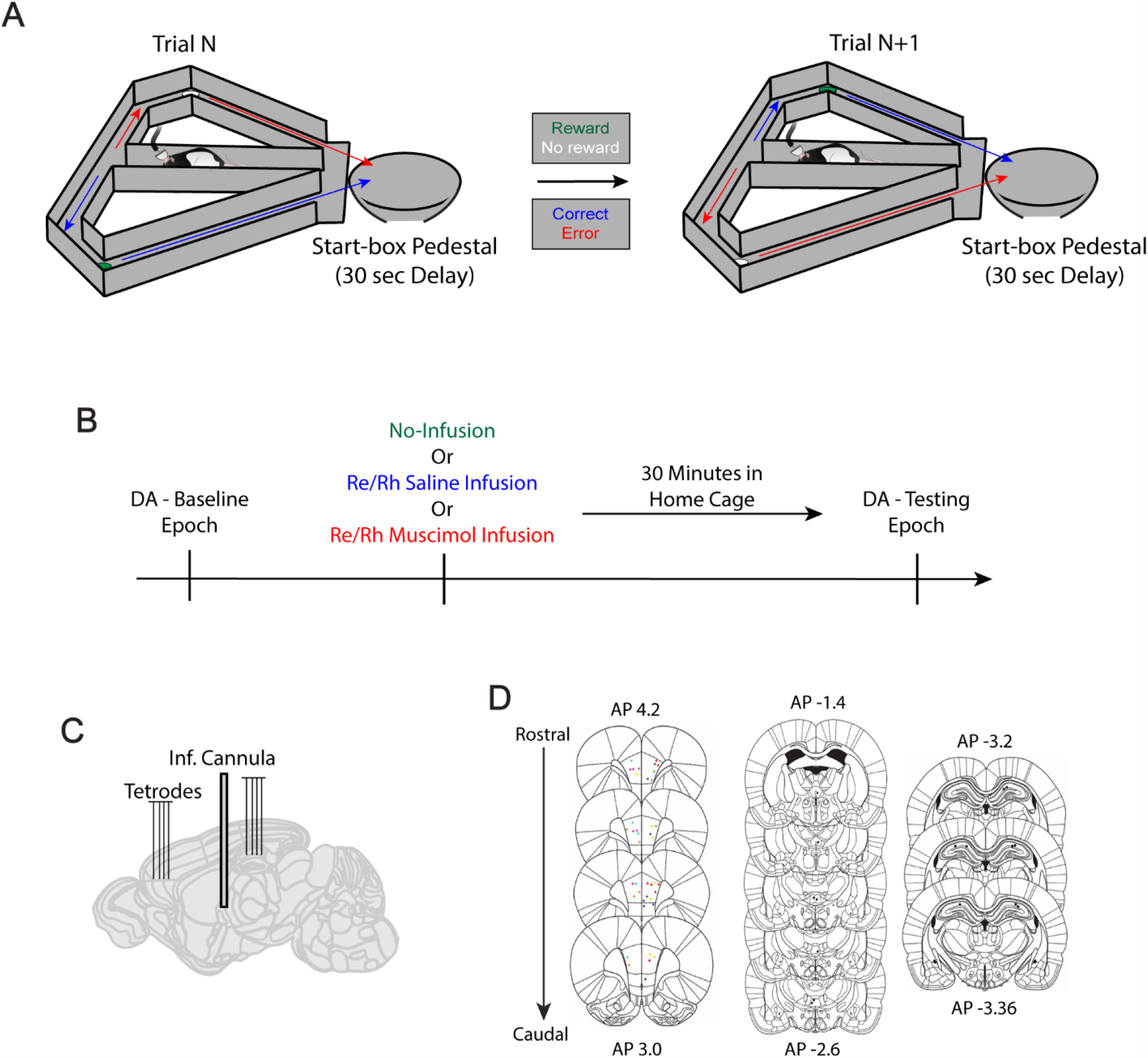
Experimental design. A) Delayed Alternation task schematic. Each trial requires the rat to alternate visits to the previously unvisited goal-arm. The first trial of the session is a free-choice trial. A 30 second delay period separates trials, during which the rat is confined to the start-box pedestal. Green cup indicates reward, white cup indicates no reward. Blue arrow indicates correct trajectory, red arrow indicates incorrect trajectory. B) During the baseline epoch, rats performed a set of 12-30 trials (DA Baseline epoch). They then received an infusion of saline, infusion of muscimol, or no infusion and returned to their home cage. Thirty minutes later, rats were tested on another set of 12-20 trials (DA Testing Epoch; See Hallock et al., 2016) C) Schematic of recording sites in the mPFC and dHPC and with a cannula site in the Re/Rh. D) Coronal stereotaxic atlas plates showing histological confirmations of tetrode and cannulae placements. Colored dots in the mPFC (left) indicate different tetrode placements (See Hallock et al., 2016).

### Handling, pre-training, and task training

Handling and pre-training were identical for data used from both studies (Hallock et al., 2013 and Hallock et al., 2016). First, rats were handled for 15 minutes per day for 5 days. They were then shaped to consume chocolate sprinkles located in plastic bottle caps at the ‘goal-zone’ of the T-maze (see **Fig. 1A** for maze schematic). Rats were given 3 minutes to consume the reward, and were then placed into the opposite goal-zone. Once rats consumed the sprinkles within 90 seconds on all 6 goal-zone visits, they were moved onto the next stage of behavioral shaping, forced runs. Prior to each forced run trial, the experimenter placed a barricade in the entry point of the left or right goal arm (6 left, 6 right trials in a pseudorandom sequence). The rats began at the start-box, then traversed the central stem of the maze, turned into the open goal arm, and were rewarded at the goal-zone. After reward consumption, rats returned to the start-box via the return arm. Once rats consumed the reward and returned to the start-box on 10/12 trials for two consecutive sessions, they began delayed alternation (DA) task training. During delayed alternation training, rats were rewarded for alternating visits to the left and right goal arms. After choosing a goal arm, the rats returned to the start box via the return arm where they were confined by a wooden barricade for 30 seconds. All rats included in the study were trained until they reached 80% proficiency on the delayed alternation task for two consecutive days (i.e. correct choices on 20/24 trials). Then they were implanted with a micro-drive loaded with independently moveable tetrodes targeting both the mPFC and the dorsal hippocampus. For Re inactivation, a guide cannula targeting the Re and the neighboring rhomboid nucleus was implanted (see Hallock et al., 2016 for surgical details).

### Recording

Each rat underwent three recording sessions separated by at least 24 hours in which one of three infusion conditions were employed: (1) no infusion (2) saline infusion into the Re, and (3) muscimol infusion into the Re (**Fig. 1B**). For each recording session, each rat performed a baseline epoch in which LFP was recorded from the dorsal hippocampus and mPFC while the rats performed a set of trials of the delayed alternation task. Immediately after the baseline epoch, the rats underwent one of the three infusions (no infusion, saline infusion, or muscimol infusion, followed 30 minutes later by another recording epoch (“testing”) during which the rat performed another set of trials in the delayed alternation task.

### Infusion Protocol

For the saline session a PBS solution (Fischer Scientific) was used. For the muscimol session a *GABA*_*A*_ receptor agonist (Life Technologies Solutions) was used. Solutions were diluted to a concentration of 0.25 μg/μL and infused using a microinfusion syringe (Hamilton) and automated infusion pump (World Precision Instruments), at a rate of 0.25 μl/min and volume of 0.5 μl.

### Perfusion and histology

Rats were given a .5 μl volume infusion of fluorophore-conjugated muscimol (BODIPY TMR-X; Life Technologies) 30 min before perfusion (Allen et al., 2008). Cannulae placements were visualized via staining half of the Re brain slices with cresyl violet, and the other half with ProLong Gold with DAPI (Life Technologies), highlighting the spread of fluorophore conjugated muscimol. Cannulae and tetrode track verifications (**Fig. 1D**) were accomplished by superimposing digital plates from the Paxinos and Watson (2006) rat brain atlas over pictures of the cresyl-stained brain slices using Adobe Illustrator.

### Behavioral analysis and VTE quantification

To quantify VTE events, we used the commonly employed integrated absolute change in angular velocity (IdPhi) method (Redish, 2016). Choice-points were visually identified as a square space surrounding the divergence in trajectory towards the goal-arms, which included position data that immediately preceded choice-point entry (see **Fig. 2A**). Position data during choice-point passes were smoothed using a gaussian-weighted moving average with a window length of 30 samples (reflects the sampling rate of ∼30 samples/sec) using MATLAB’s *smoothdata* function. IdPhi was calculated using LED tracking of the rats’ head-stage (Papale et al., 2012; MATLAB code provided by David Redish). Briefly, for 2-dimensional position data, velocity in the × (‘dX’) and Y (‘dY’) dimensions were obtained using a discrete time-adaptive windowing method (Janabi-Sharifi et al., 2000). Phi was then defined by taking the atan2 of dY and dX. Next, the change in movement orientation (dPhi) was estimated by applying the time-adaptive windowing method to the unwrapped phi estimate. The integrated absolute value of change in movement orientation (IdPhi) was defined by taking the integral of |dPhi| estimates. Finally, the natural log and z-score of all IdPhi scores was taken to produce the zlnIdPhi distribution (**Fig. 2A**).

**Figure 2:**
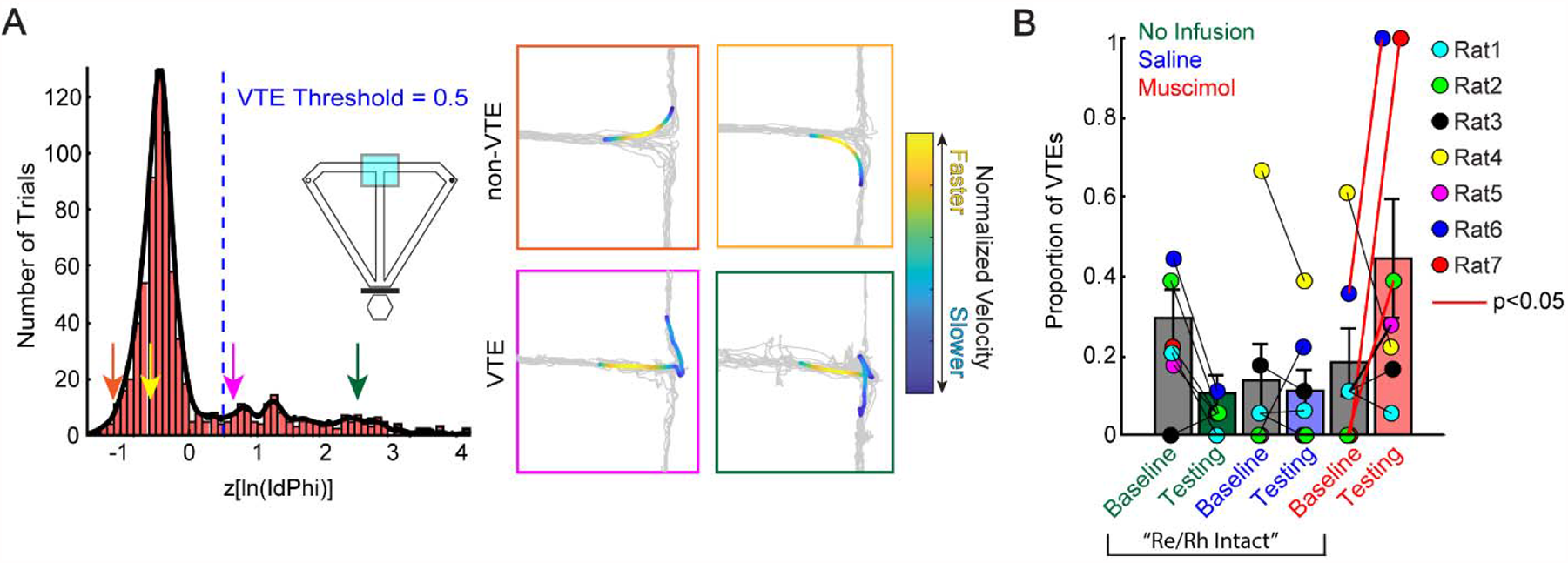
Re inactivation has variable effects on the frequency of VTEs. A) zlnIdPhi distribution. VTE threshold was set to 0.5, similar to previous reports (see methods). Arrows are color-coded to denote example VTE and non-VTE passes in the right panel. Notice that VTEs exhibit complex head-movement behaviors compared to stereotypical, or ballistic-style, non-VTEs. B) Re inactivation variably increased the frequency of VTE behaviors. Colored dots indicate individual rats. There was a reduction in VTE proportion from baseline to testing in the no infusion condition that did not survive alpha correction. There was no significant change from baseline to testing in the saline condition or in the muscimol condition. Data are displayed as the mean ± SEM. Note Re inactivation led to an increase in VTE behaviors in some rats and not others. Specifically, rats 2, 6 and 7 showed a significant increase in zlnIdPhi under Re suppression. Red solid lines indicate a significant increase in the zlnIdPhi score using a two-sample t-test with trials as the sample-size.

The VTE threshold was estimated by first identifying the two prominent ‘components’ of the distribution the ‘normal’ and ‘tail’ components (**Fig. 2A**, see Redish, 2016). The ‘normal’ component reflects ballistic-like trajectories (see **Fig. 2A**), and were the predominant trajectory type across trials. To extract the ‘tail’ component that reflects VTE behaviors, we used 0.5 standard deviations of the lnIdPhi score as a threshold. This cut-off is highly consistent with past work (Papale et al., 2012; Redish, 2016; Amemiya and Redish, 2018) and was visually confirmed to contain pause-and-reorient behaviors (**Fig. 2A**). Behavior during VTE trials included head-sweeps, looking around for an extended period at the T-intersection, and prolonged pauses before making a decision. Code for this analysis can be found at www.github.com/GriffinLabCode/GriffinCode.

In spatial alternation tasks, perseveration behaviors are reflected by repeated same-turn decisions and reflect behavioral inflexibility (Viena et al., 2018). To assess perseveration strategies, we used a perseveration index. This index was defined by dividing the sum of 2 consecutive errors (or 3 same direction turns) by the amount of possible perseveration instances, similar to a previous publication from our lab (**Fig. 3A;** Hallock et al., 2013).

**Figure 3:**
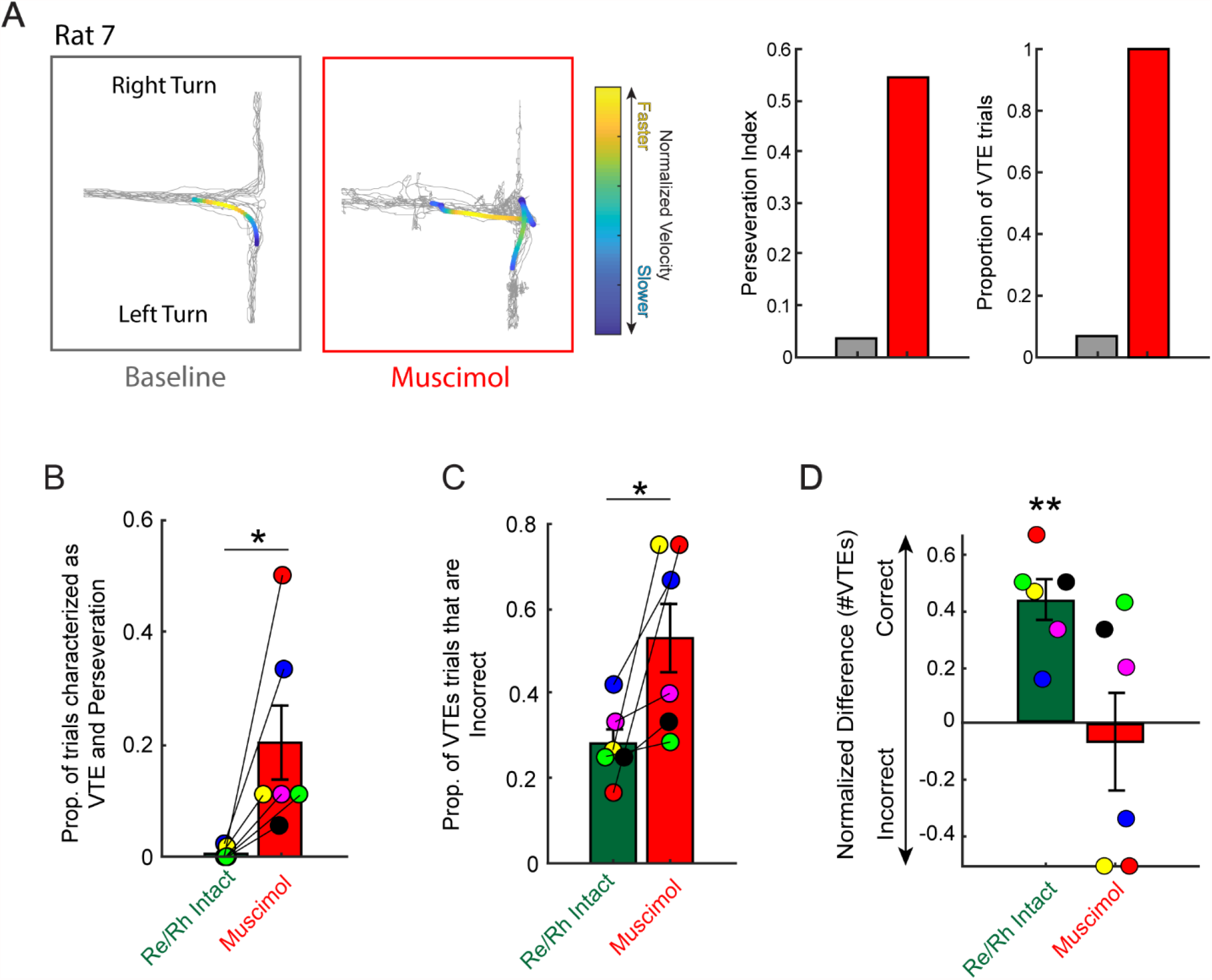
Re inactivation impacts the *success* of VTEs. A) An example rat that showed both high perseveration (multiple same direction errors) and a large number of VTEs after Re suppression. B) Re inactivation was associated with a significant increase in co-occurring VTE and perseverations (back-to-back errors). Data are displayed as the mean ± SEM. C) Re inactivation led to an increase proportion of VTEs that occurred during an erroneous choice (i.e. #VTEs on an error trial / total #VTEs). D) Normalized difference of #VTEs ([#VTE correct - #VTE incorrect] / [#VTE correct + #VTE incorrect]) revealed that when the Re was intact, VTEs were often associated with a correct decision. Under Re inactivation, VTEs were not significantly associated with correct or incorrect decision-making. There was a pre-alpha correction significant difference between Re intact and muscimol groups. Data are presented as the mean ± SEM. Colored dots indicate individual rats as in **Fig. 2**. ^*^p<0.05, ^**^p<0.01.

### Spectral analysis

Spectral analyses were performed using the chronux toolbox in MATLAB (Mitra, 2007). The entire recorded LFP signal was first detrended using *locdetrend*, then scrubbed of noise using *rmlinesmovingwinc*. A window length of 1-second with a 500ms overlap was used for detrending and denoising. 5 multi-tapers with a time-bandwidth product of 3 were used for analysis. Theta was defined as 5-10 Hz based on observations of the mPFC and HPC power spectra. Clipping artifacts were detected using *detect_clipping*.*m* (an in-house algorithm that can be found on the lab’s github page) and if the number of artifacts exceeded 1% of the signal length, the trial was excluded. To detect clipping events, we took the derivative of the LFP signal to find repeating LFP estimates. Using this output vector, we again performed the derivative to extract multiple repeating elements, which we defined as clipping artifacts. Visual inspection of data (when sampled at ∼2034Hz) with clipping artifacts revealed that this method adequately extracted these events from the dataset. Rats contributing fewer than 3 trials were excluded from analysis. In the case where mPFC and HPC LFP were sampled at different sampling frequencies, the longer LFP segment was down-sampled and the shorter segment was interpolated to match the updated long segment. After down-sampling, LFP segments were often different by a single sample and therefore this interpolation did not change features of the LFP.

### Statistical analysis and figure generation

To correct for multiple comparisons, Bonferroni’s correction was applied. Statistical procedures and figure generation were performed using custom MATLAB code and IBM SPSS. Adobe Illustrator was used for figure creation.

## Results

### Rats engage in varying strategies following Re inactivation

Rats were trained to perform a delayed alternation task and then implanted with microdrives targeting the mPFC and dHPC along with an infusion cannula targeting the Re (see methods, **Fig. 1C and 1D**). For each recording session, rats first performed a set of delayed alternation trials (“DA Baseline Epoch”). Next, they underwent one of three experimental manipulations: no-infusion, saline-infusion, or muscimol-infusion into the Re. After a 30-minute waiting period in their home cage, rats performed a second session of delayed alternation (“DA Testing Epoch”; **Fig. 1B**). Electrophysiological recordings were taken from the mPFC and dHPC during both baseline and testing task epochs (**Figs 1C and 1D**).

We first examined the impact of Re inactivation on VTE behaviors. Trials were excluded from the VTE analysis if they exhibited obvious signs of video-tracking errors that would interfere with IdPhi estimation. We excluded data from the no infusion condition of one rat due to numerous tracking errors on the no-infusion baseline session. We found that Re inactivation did not reliably increase VTE behaviors across all rats under muscimol suppression (**Fig. 2B;** t(6) = -1.48, p = 0.19, α = 0.0167), nor after infusion of saline (t(6) = 0.436, p = 0.68, α = 0.0167). Interestingly, however, there was a reduction in VTE frequency between no infusion baseline and no infusion testing that did not survive alpha correction (t(5) = 3.09, p = 0.027, ci = [.031 .338], α = 0.0167). This reduction in the occurrence of VTEs was likely due to repeated exposure to the task, suggesting a shift in learning stages from deliberation to planning (Hu & Amsel 1995; Bett et. al., 2012; Papale et. al., 2012; Schmidt et al., 2013; Redish, 2016). Due to notable numerical increases in VTE frequency from baseline to muscimol testing in some rats, we were curious as to how many rats showed a significant increase in VTE behaviors with Re inactivation. Therefore, for each rat, we performed two-sample t-tests between baseline and testing zlnIdPhi scores for each condition (no infusion, saline, muscimol). After Bonferroni alpha corrections, 3/7 rats showed a significant increase in zlnIdPhi with Re inactivation (**Fig. 2B**, Rat7: t(30) = 24.57, p<.001; Rat6: t(24) = 4.37, p<.001; Rat2: t(34) = 3.25, p<0.01; alpha set to 0.0167). Thus, fewer than half of the rats exhibited more deliberative behaviors under Re suppression. In effect, these results indicate that Re inactivation does not *cause* VTE behaviors, but that some rats reverted to deliberation when the Re was disrupted.

### Re inactivation reduces the success of VTEs

Because VTEs are thought to reflect periods of deliberation, VTEs should help the rat choose the correct goal arm during the delayed alternation task. Yet, when visualizing choice-point behavior during muscimol testing sessions, we observed one rat that exhibited VTEs on every trial, had a profound turn-bias, and performed only 31% of trials correctly (**Fig. 3A**, notice the density of left trajectories compared to right). Observations from 2 other rats in the dataset revealed similar behavior, namely, the co-occurrence of VTEs and perseverative errors (i.e. repeated, same-sided choices; see methods). Thus, we next wondered if deliberation and perseverative behaviors reliably overlapped when the Re was inactivated. To examine this question, we estimated the proportion of trials that were categorized as both a VTE and a perseverative error. Since VTEs are primarily seen during learning (Redish, 2016), and all subjects included in the study were trained to task-proficiency, we were not surprised to obtain small samples of these behaviors within each control condition (all conditions except muscimol testing). Therefore, to maximize the chance that we extracted correct and incorrect VTE behaviors during well-learned behavior, we collapsed across control conditions per each rat, generating a single control variable that contained every trial except those in the muscimol session (*Per each rat*, #*VTE trials in control group*: 6, 19, 6, 30, 8, 8, 10; *#VTE trials in muscimol group*: 12, 12, 5, 4, 3, 7, 1). For purposes of clarity, we refer to this variable as ‘Re intact’. We found that Re inactivation was associated with a significant increase in VTEs that occurred during a perseverative behavior (**Fig. 3B;** t(5) = -2.78, p = 0.039, ci = [-.379 -.015], cohens D = 1.59, α = 0.05). This finding directly corroborates our visual observations of co-occurring perseveration and VTE behaviors under Re suppression (**Fig. 3A**). We then investigated whether or not Re inactivation led to an increase incidence in VTE behaviors during non-perseverative errors. Similar to our results in **Fig. 3B**, Re inactivation led to a significant increase the amount of VTEs that occurred on error trials (**Fig. 3C**; t(5) = -2.62, p = .047, ci = [-.495 -.005], cohens D = 1.53, α = 0.05). Thus, Re suppression was associated with an increase in deliberative behaviors that result in an incorrect choice.

Since VTE behaviors reflect an early learning strategy, decrease with task proficiency (Papale et al., 2012), and are thought to be a behavioral manifestation of deliberation (Redish, 2016), VTEs should help decision-making when observed during well-learned performance. To test whether VTEs were associated with a greater chance of observing a correct choice, we compared the amount of correct and incorrect VTE behaviors under Re intact and Re suppression conditions. Consistent with our results in **Fig. 3A-C**, we found that VTEs were only significantly associated with correct decision-making when the Re was intact (**Fig. 3D**; Re Intact: t(5) = 6.18, p = 0.002, ci = [.256 .619], cohens D = .97, α = 0.0167; Muscimol: t(5) = -.35, p = .74, α = 0.0167; t-tests against a null of 0). There was a pre-alpha level correction difference between Re intact and Muscimol conditions (t(5) = 2.62, p = .047, ci = [.009 .99], α = 0.0167, paired t-test). It should be noted, however, that a caveat to this analysis is that there are more correct than incorrect trials in the ‘Re intact’ dataset. Moreover, it should be noted that there were generally more errors under Re suppression; however, it is interesting nonetheless, that rats exhibited simultaneous perseverative errors with VTE behaviors. Taken together, our results suggest that VTEs are associated with correct decision-making in the intact brain and in well-trained animals. However, with Re inactivation, the incidence of errors increases despite the occurrence of VTE behaviors. Thus, Re inactivation reduces the ‘success’ of deliberative-behaviors.

### Re suppression induces behavioral inflexibility patterns observed during early-learning

Thus far, we have demonstrated that Re suppression results in variable frequency of VTEs, but when those VTEs occur, they do not reliably help rats make a correct decision. Given the poor choice accuracy observed under Re suppression (Hallock et al., 2016) along with failed VTEs, it was next important to characterize the types of errors that we observed. For example, repetitive turning into one of the goal arms (perseveration, see **Fig. 4A**) would suggest behavioral inflexibility, a finding that we visually observed in our data (**Fig. 3A and Fig. 4B**), and that has been reported on a different place-memory task (Viena et al., 2018). However, it was additionally important to characterize other kinds of errors. For example, it was possible that rats were using an error-feedback strategy, by using an incorrect choice to attempt to make a correct decision on the next choice-point pass (**Fig. 4A**).

**Figure 4:**
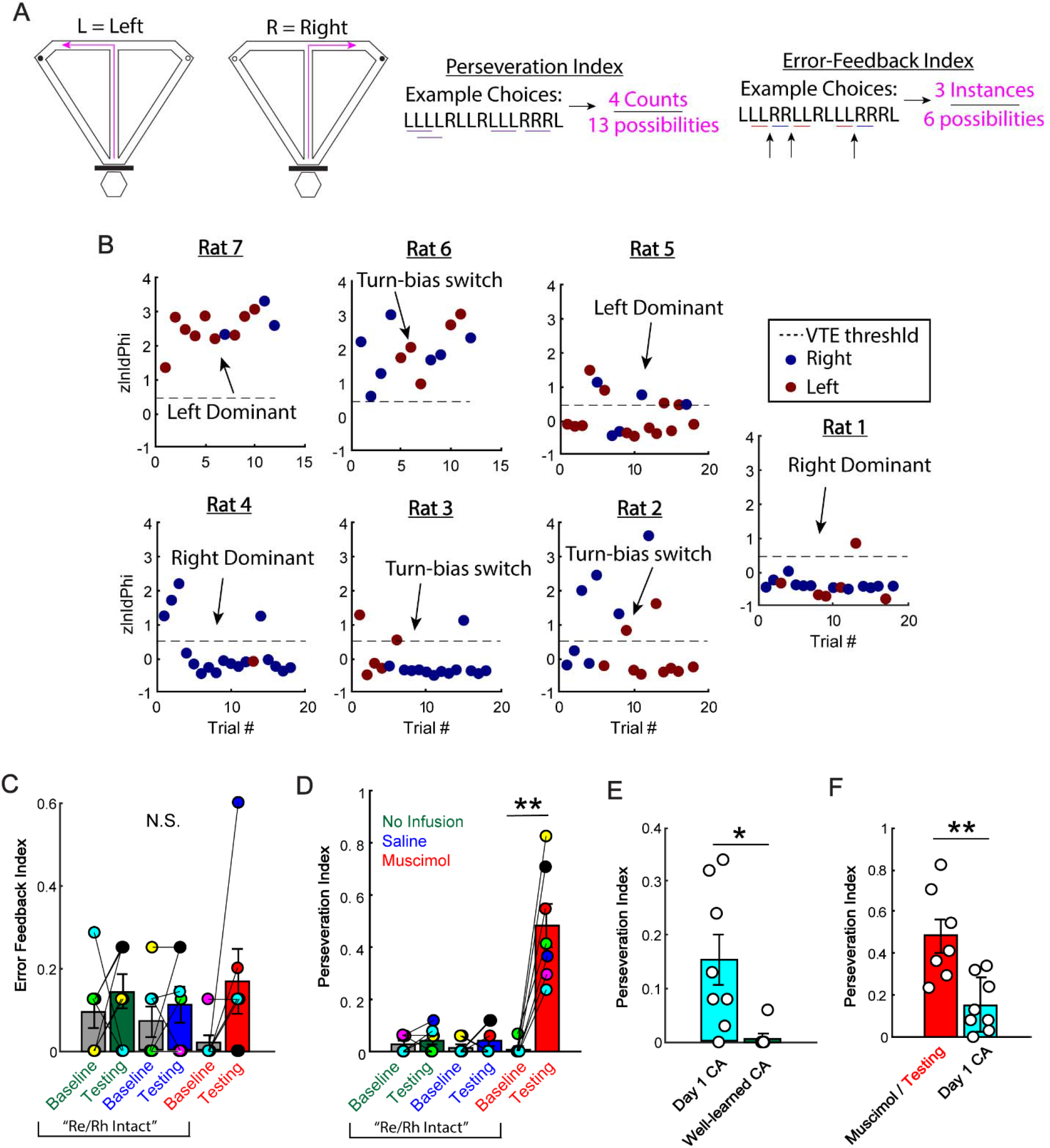
Re suppression induces inflexible decision-making strategies normally observed during early learning. **A)** Schematic of the DA task with examples of possible patterns of decision-errors. A perseveration index (# back-to-back errors / # possible back-to-back errors) and error-feedback index (# instances where an error in one direction results in an error in the other direction / # possible instances). **B)** Scatter plot of zlnIdPhi scores across trials for each rat. Blue dots indicate a right choice, red dots indicate a left choice. Notice the clear examples of perseveration with VTEs being variably applied. **C)** Muscimol suppression resulted in no significant change in error-feedback strategies across conditions. Data are displayed as the mean ± SEM. **D)** Muscimol suppression resulted in a drastic increase in same-turn decisions, which was quantified using a perseveration index. **E)** Similar to (C), except on day 1 learning of CA and well-learned CA performance in intact rats. Notice that turn-bias behaviors are present during initial learning, and reduce when the same rats perform above 80% for at least two consecutive days. ^*^p<0.05. **F)** Re inactivation increased perseverative behaviors beyond what was seen during day 1 performance of the CA task in intact rats. ^**^p<0.01. White circles indicate individual rats. Data from CA rats were taken from Hallock et al., 2013, while data from DA rats were taken from Hallock et al., 2016.

When visualizing behavioral patterns across rats, it was clear that rats exhibited various strategies when the Re was inactivated. While some rats deliberated throughout the entirety of the session (**Fig. 4B**), every rat exhibited dominant perseverative behaviors similar to what has previously been reported (Viena et al., 2018). Additionally, many rats exhibited clear preferences in turn direction (**Fig. 4B**). Interestingly, across animals, perseverative behaviors were typically followed by an alternation or turn-bias ‘switch’ at some point in the session (**Fig. 4B**). For example, the turn choice shown in the second column of **Fig. 4B** exhibited the following pattern; R-R-R-R (switch) L-L-L (switch) R-R. This observation indicates that some rats were attempting to switch decision-strategies, but remained largely inflexible in behavioral adaptation. In line with this idea, some rats seemed to deliberate sparsely throughout the session (**Fig. 4B**). However, it was clear that a perseverative strategy (**Fig. 4D**; Baseline: t(5) = -.6 p = .57; Saline: t(6) = -.89, p = .407; Muscimol: t(6) = -5.61, p = .001, CI = [-0.68 -0.27], cohens D = 3.0, α = 0.0167, paired t-tests), but not an error-feedback strategy (**Fig. 4C;** No infusion: t(6) = -0.69 p = 0.515; Saline: t(6) = -.857, p = 0.464; Muscimol: t(6) = -1.86, p = 0.11; paired t-tests) dominated the choice process. These results support the conclusion that Re inactivation causes behavioral inflexibility (Viena et al., 2018).

In place memory tasks, deliberation has been theorized to reflect the first stage of learning where rats understand their environment, but not the rules to attain a reward (Redish, 2016). In contrast to the utility of VTE behaviors, perseverative strategies are maladaptive to alternation tasks, and likely reflect competing habit-based processes. This maladaptive strategy was clearly demonstrated by (Packard et al., 1989), showing that hippocampal lesions facilitated the learning of a striatal-dependent task. However, recent evidence indicates that the addition of turn-biases improved the accuracy of behavioral models on predicting spatial alternation learning (Kastner et al., 2020). Because we observed variability in the application of VTE behaviors (**Fig. 2**), it was possible that Re suppression reverted rats to a state of learning. The logic of this idea holds that rats would variably express VTE’s as a strategy to access HPC-dependent information (Redish, 2016). However, by disconnecting the mPFC from the dHPC, this process would fail, resulting in increased errors that co-occur with VTEs (**Fig. 3**). Thus, if Re inactivation reverted rats to a state of learning, but a competing brain-system dominated the decision-process, then the perseverative strategy should be apparent on initial exposure to the alternation rule.

To test this idea, we next quantified the types of errors occurring on day 1 of learning the continuous alternation (CA) task (Hallock et al., 2013). This task is identical to delayed alternation except that there is no delay imposed between trials. Thus, rats are required to learn the spatial alternation rule, without having to retain trial-specific information over a delay period. We found that on the initial exposure to the alternation rule (i.e. day 1 of learning of the CA task), rats exhibited greater perseverative behaviors when compared to asymptotic performance (**Fig. 4E;** t(7) = 3.28, p = .013, CI = [.040 .245], cohens D = -1.58, α = 0.05, paired t-test). We then compared Re inactivated behavior to that of day 1 CA learning, and found an exaggeration of perseverative strategies in the Re inactivated group when compared to day 1 of learning a CA task (**Fig. 4F;** t(13) = 3.69, p = .003, CI = [.139 .530], cohens D = -1.87, two-sample t-test). These results support the idea that Re inactivation led to a competing brain-system dominating the decision-process, as the choice-errors reflect early-learning behaviors. Moreover, because VTE behaviors were variably applied, these result further support the idea that some rats attempted to re-learn the delayed alternation task, but place-memory utility was prevented by disconnecting the mPFC and HPC through Re suppression.

### Re suppression reduces prefrontal-hippocampal theta coherence during VTEs

We next set out to address the hypothesis that Re orchestrates dHPC-mPFC synchrony during VTE behaviors specifically by examining the impact of Re inactivation on mPFC-dHPC coherence during VTE behaviors. First, we extracted and cleaned LFP from the choice-point (see methods). We then calculated coherence, down-sampled spectral estimates to match the lowest Nyquist frequency, then examined the coherence distribution. Similar to our visual observations in the power spectra that drove our definition of theta (5-10 Hz, see methods), there was a peak in the coherence distribution at the theta range. We then extracted and averaged across those frequencies and found that Re inactivation significantly reduced theta-coherence during trials in which a VTE was observed (VTE trials) (**Fig. 5B;** t(5) = 3.1, p = 0.027, CI = [.016 .169], cohens D = -1.47, paired t-test). Interestingly, however, we did not find a reduction in theta coherence during trials in which a VTE was not observed (non-VTE trials) (**Fig. 5C;** t(4) = 0.472, p = 0.66, paired t-test). Thus, in well-trained animals, these findings indicate that Re inactivation impacts mPFC-dHPC theta coherence specifically during VTE behaviors.

**Figure 5:**
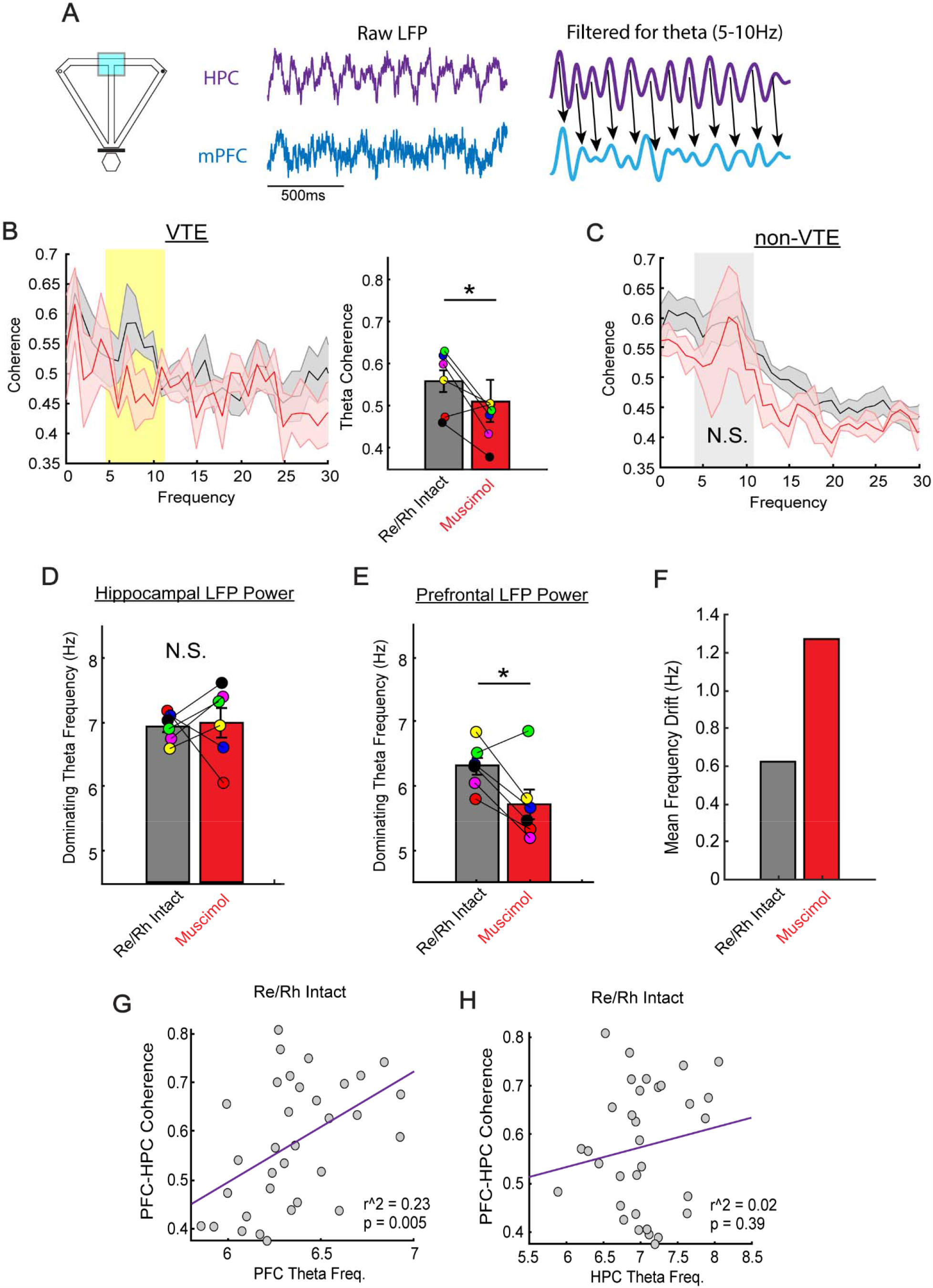
Re inactivation reduced prefrontal-hippocampal theta coherence during VTEs through its contribution to the prefrontal theta oscillation. A) Schematic of the maze illustrating the region of the maze occupied by the rat for the theta coherence analysis and example LFP. The right panel demonstrates a visualization of coherence, a metric of spectral correlation, with black arrows illustrating the consistency of the peak-to-peak correspondence in the two signals. B) Frequency × coherence distribution from Re intact (grey) and muscimol (red) data. Notice the increased coherence in the theta range (5-10Hz) as demarcated by a yellow box. During VTE trials, there was a specific reduction in theta (5-10Hz) prefrontal-hippocampal coherence. *Right panel* demonstrates that Re inactivation significantly reduced theta-coherence. ^*^p<0.05. Data are displayed as the mean ± SEM. C) On non-VTE trials, there was no significant reduction in theta coherence. The grey box demarcates the theta range. D) Re inactivation did not significantly impact the peak theta frequency in the dHPC during VTE trials. Note however, some individual rats showed an increase in theta frequency after Re inactivation. E) Re inactivation significantly reduced the peak theta frequency in the mPFC during VTE trials. ^*^p<0.05 F) Mean frequency shift was defined by the difference in rat-average coherence scores. As seen by comparing (D to E), the dominating theta frequency in the mPFC and dHPC, during Re inactivated VTEs, was further apart in comparison to when the Re was intact. G) mPFC-dHPC theta coherence is strongly predicted by changes in the dominating theta frequency in the mPFC. N = 33 control sessions across 7 rats. H) mPFC-dHPC theta coherence is not strongly predicted by changes in the dominating theta frequency in the HPC. N = 33 control sessions across 7 rats.

Since coherence is estimated by dividing the cross-spectral power by the sum of both signals’ power spectra, it was important to identify *how* theta coherence was being modulated by Re inactivation. Therefore, we next examined the impact of Re inactivation on peak theta frequency, the theta frequency that exhibited the strongest spectral power, in both the dHPC and mPFC. To account for the effect of the 1/f power law, we logarithmically transformed the data before extracting the peak theta frequency. We found that Re inactivation significantly reduced the mPFC peak theta frequency (**Fig. 5E**; t(5) = 2.93, p = 0.033, CI = [.072 1.12], cohens D = -1.19, paired t-test), but did not impact the dHPC peak theta frequency (**Fig. 5D;** t(5) = -0.23, p = 0.829, paired t-test). Thus, Re inactivation reduced mPFC-dHPC theta coherence by shifting the mPFC theta oscillation further from the dHPC theta oscillation by an average of > 1.2Hz (**Fig. 5F**). These findings indicate that Re inactivation primarily affected mPFC-dHPC theta coherence by directly modulating the mPFC theta oscillation.

Next, it was important to characterize whether mPFC theta normally drives variability in mPFC-dHPC theta coherence. Similar to previous work (O’Neill et al., 2013), we reasoned that statistically removing the influence of dHPC theta on mPFC-dHPC theta coherence could provide insight into causal influence of each region on interregional theta synchronization. Therefore, we performed a multiple regression on the LFP data from ‘Re intact’ sessions using mPFC and dHPC peak theta frequency as predictors. After controlling for the influence of dHPC theta frequency on mPFC-dHPC theta coherence, mPFC theta frequency was a strong predictor of variability on coherence (zero order R = 0.48, part R = 0.455, t(32) = 2.8, p = 0.008). However, dHPC theta frequency was not a strong predictor of mPFC-dHPC theta coherence regardless of whether the influence of mPFC theta frequency was being controlled (zero order R = 0.16, part R = -0.021, t(32) = -.132, p = 0.896). Thus, variability in how similar the mPFC theta is to dHPC theta strongly accounts for mPFC-dHPC theta coherence.

Finally, these analyses required us to address the influence of overt behavior (i.e. time-spent/running speed) on our measures of spectral analysis. To assess the influence of time-spent on estimates of theta coherence, we performed multiple analyses. First, we examined if there were statistical differences in time-spent during VTE behaviors (between Re intact and muscimol data). While there was no significant difference in time-spent between Re intact and muscimol suppression conditions, 5/6 rats showed numerical increases in time-spent during VTEs (**Fig. 6A;** t(5) = -1.5, p = 0.19, ci = [-8.7 2.2], paired t-test). Therefore, next we performed correlations between time-spent at the choice-point and coherence estimates for each session, and found no significant correlation (**Fig. 6B**; R = -0.06, p = 0.73). Thus, variability in time-spent between groups does not explain differences in theta coherence. Importantly, time-spent was also not significantly correlated with the peak theta frequency in the mPFC (**Fig. 6C;** R = -0.25, p = 0.12). However, as would be expected (McFarland, 1975; Sławińska and Kasicki, 1998; Hinman et al., 2011; Bender et al., 2015), the peak theta frequency in the HPC was negatively correlated with time-spent (**Fig. 6D;** R = -0.35, p = 0.028), and therefore positively associated with running velocity (**Fig. 6E**; R = 0.34, p = 0.032), although the relationship to velocity is likely indirect and through changes to running acceleration (Kropff et al., 2021).

**Figure 6:**
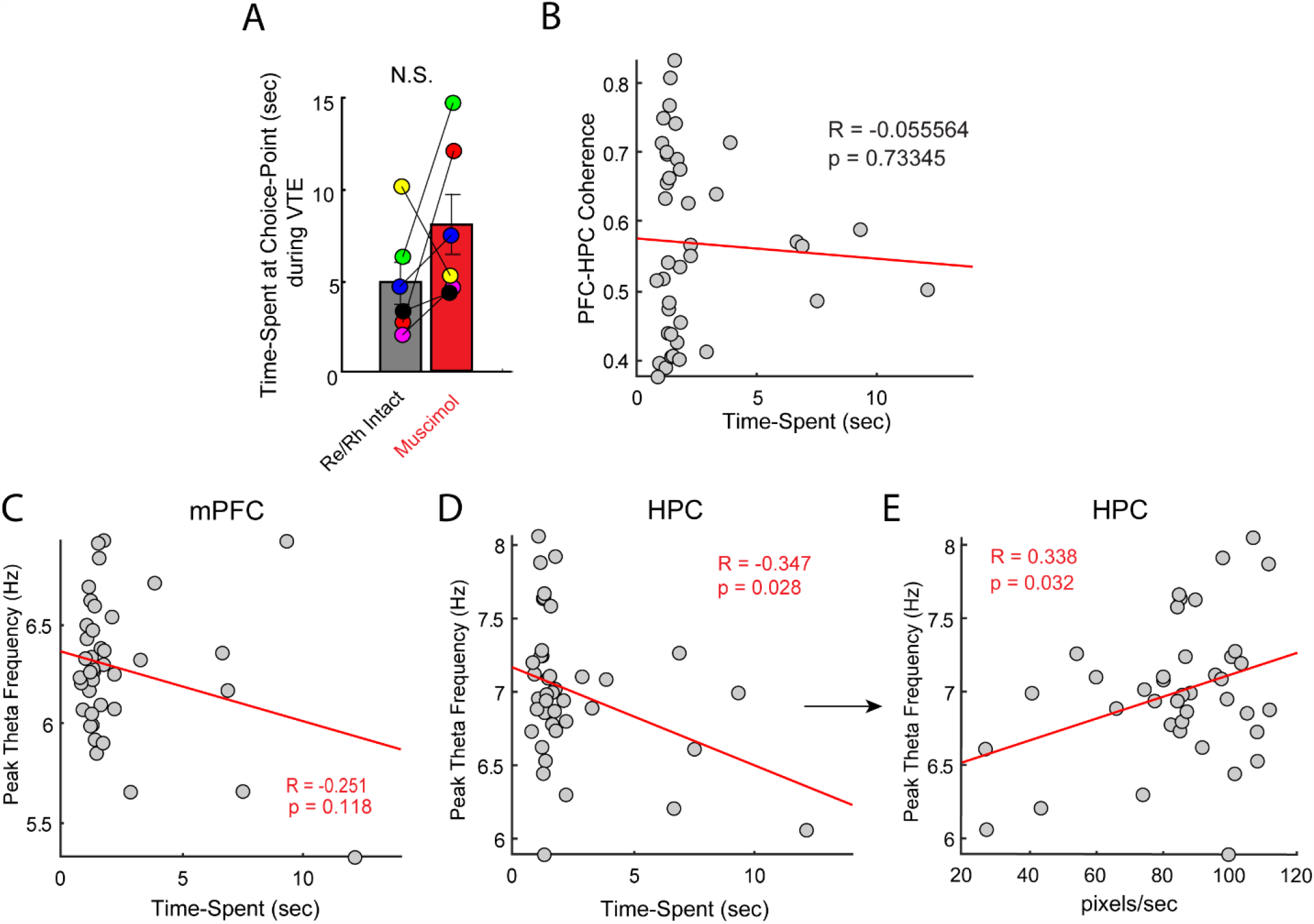
Relationship between time-spent at the choice-point and spectral analysis. **A)** There was no significant difference between the amount of time-spent at the choice-point during VTEs, between muscimol and Re intact data. Note that 5/6 rats showed numerical increases in time-spent during VTEs under Re suppression. **B)** There was no significant correlation between time-spent at the choice-point and mPFC-HPC theta coherence. Each dot is a session with coherence and time-spent estimates averaged across all trials. **C)** The peak theta frequency in the mPFC was not significantly correlated with time-spent at the choice-point. Dots indicate sessions. **D)** The peak theta frequency in the HPC was negatively correlated with the amount of time-spent at the choice-point. **E)** Similar to **(E)**, the peak-theta frequency in the HPC positively scales with running speed at the choice-point.

Finally, we statistically controlled for overt behavior by performing multiple regression similar to the results shown in **Fig. 6**. Thus, the following variables were treated as predictors of mPFC-dHPC theta coherence: mPFC peak theta frequency, dHPC peak theta frequency, time-spent at choice-point, running velocity at the choice-point, and zlnIdPhi at the choice-point. All sessions were used in this analysis since Re suppression impacted mPFC peak theta frequency and numerically increased the time-spent at the choice-point in 5/6 animals. Despite controlling for both dHPC peak theta frequency and overt behavior on mPFC-dHPC theta coherence, mPFC peak theta frequency remained the strongest predictor of mPFC-dHPC theta coherence (**Table 1a and Table 1b**). Interestingly, running velocity was another, albeit weaker, predictor of mPFC-dHPC theta coherence.

**Table 1a:** Multiple regression on mPFC-dHPC theta coherence.

| Model |  | Unstandardized Coefficients |  | Standardized Coefficients | t | Sig. |
| --- | --- | --- | --- | --- | --- | --- |
|  |  | B | Std. Error | Beta |  |  |
| 1 | (Constant) | -1.176 | .393 |  | -2.989 | .005 |
|  | PFC Peak Theta | .214 | .054 | .574 | 3.929 | .000 |
|  | HPC Peak Theta | .006 | .043 | .022 | .138 | .891 |
|  | Time-Spent | -.004 | .024 | -.074 | -.163 | .872 |
|  | Velocity | .004 | .002 | .733 | 2.439 | .020 |
|  | zlnldPhi | .157 | .095 | .850 | 1.652 | .108 |

a. Dependent Variable: mPFC-dHPC Theta Coherence

**Table 1b:**
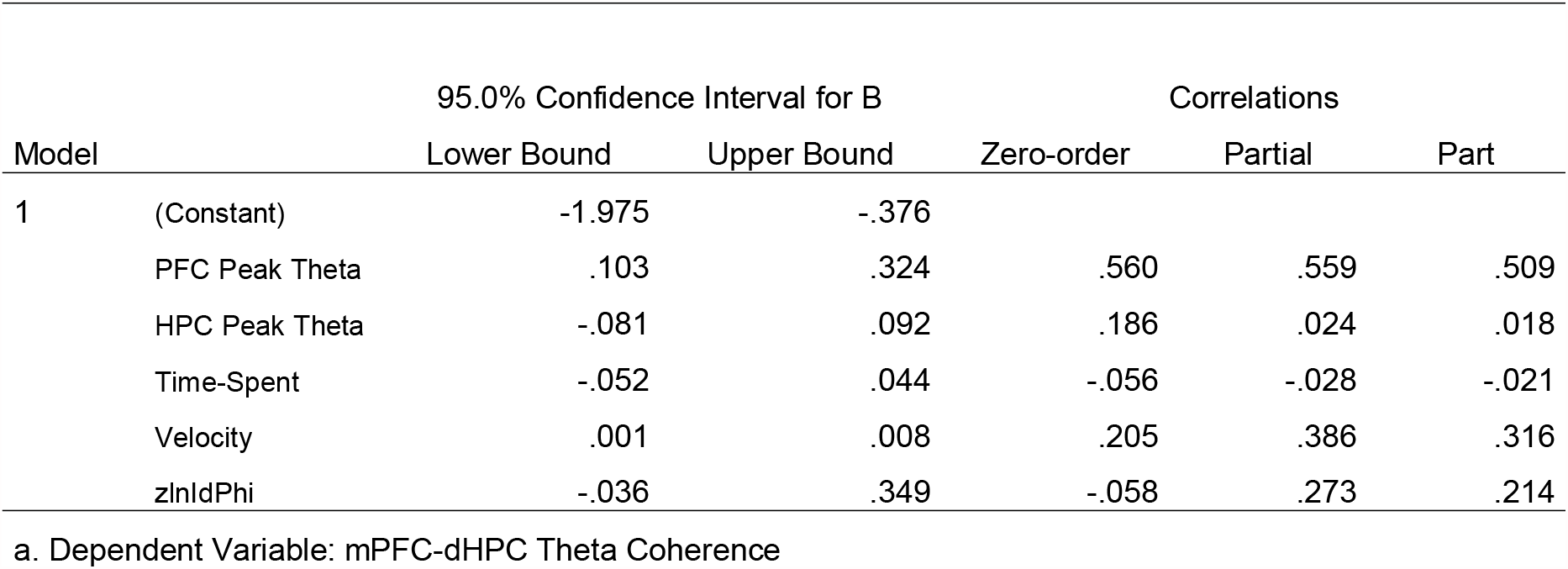
mPFC peak theta power remains a strong correlate of mPFC-dHPC theta coherence After accounting for multiple behavioral variables with multiple regression.

| Model |  | 95.0% Confidence Interval for B |  | Correlations |  |  |
| --- | --- | --- | --- | --- | --- | --- |
|  |  | Lower Bound | Upper Bound | Zero-order | Partial | Part |
| 1 | (Constant) | -1.975 | -.376 |  |  |  |
|  | PFC Peak Theta | .103 | .324 | .560 | .559 | .509 |
|  | HPC Peak Theta | -.081 | .092 | .186 | .024 | .018 |
|  | Time-Spent | -.052 | .044 | -.056 | -.028 | -.021 |
|  | Velocity | .001 | .008 | .205 | .386 | .316 |
|  | zlnldPhi | -.036 | .349 | -.058 | .273 | .214 |

a. Dependent Variable: mPFC-dHPC Theta Coherence

Taken together, our results indicate that Re inactivation reduces mPFC-dHPC theta coherence during VTE trials. Moreover, our findings indicate that the Re contributes to mPFC-dHPC theta coherence through its influence on the mPFC theta oscillation. Finally, our analysis suggest that mPFC-dHPC theta coherence is predominately driven by variability in the mPFC theta oscillation during choice-point behaviors during asymptotic performance of a spatial-working memory task.

## Discussion

In this report, we examined the impact of Re inactivation on deliberative behaviors. We predicted that an intact ventral midline thalamus would be important for the behavioral manifestation of VTE behaviors. Instead, we found that Re inactivation increased deliberative errors -- choice-errors that occurred simultaneously with a VTE. Moreover, these data highlight a strong contribution of the Re on the prefrontal theta oscillation because Re inactivation reduced the mPFC peak theta frequency during deliberation, effectively de-synchronizing prefrontal and hippocampal theta oscillations. Detailed examination of choice behaviors indicated that under Re suppression, rats reverted to maladaptive strategy marked by perseverative errors and the variable application of a deliberative strategy. Our work provides the first demonstration of how the ventral midline thalamus contributes to mPFC-HPC communication during deliberation. Importantly, this study highlights a specific contribution of the Re on mPFC neural oscillations during deliberation.

During the learning of T-maze paradigms with difficult choices, it has been hypothesized that rodents undergo three stages of learning; deliberation, planning, and automation (Redish, 2016). During the deliberation stage, VTEs exist at high-cost choice-points, and decrease with mPFC or dHPC inactivation (Hu and Amsel 1995; Bett et. al., 2012; Schmidt et. al., 2019; Kidder et al., 2021). By inactivating the Re, we experimentally disconnected the mPFC from the dHPC which reduced the ability for VTEs to ‘help’ rats make a correct choice. Thus, our findings suggest that the Re may coordinate critical information for *successful* deliberation, such as the communication of potential outcomes computed by the HPC (Johnson and Redish, 2007) or goal-relevant information from the mPFC (Hasz and Redish 2020). In support of this idea, mPFC-dHPC theta coherence, a measure of neural synchronization, was reduced during VTEs under Re suppression. This reduction in mPFC-dHPC theta coherence appears to be driven by a reduction in the peak theta frequency in the local mPFC theta oscillation. Importantly, these findings are in agreeance with recent work suggesting that mPFC-dHPC theta coherence is linked to epochs of elevated attention (Bygrave et al., 2019), as would be required during deliberation.

It has been demonstrated that as rats improve choice-accuracy on spatial memory tasks, mPFC-dHPC theta coherence increases (Benchenane et. al., 2010; Hallock et. al., 2016). However, the opposite is true of VTE behaviors, such that they become less apparent with proficient learning (Papale et. al., 2012; Redish, 2016). While VTEs were not a significant predictor of mPFC-dHPC theta coherence, it is important to note that all recordings were done in well-trained rats. Therefore, we cannot not rule out the possibility of a relationship between VTEs and mPFC-dHPC theta coherence during task acquisition. However, our linear regression analyses indicated that variability in the dominating mPFC theta frequency was driving changes to mPFC-dHPC theta coherence. Thus, a prediction that arises from these analyses is that mPFC-dHPC theta coherence increases with learning as decision-systems (mPFC) learns to ‘tune in’ to spatial-memory systems (dHPC), but not the other way around. If true, we would expect that with learning, a negative relationship should exist between VTE occurrence and the dominating mPFC theta frequency. This finding would indicate that as rats deliberate less, and therefore make their decision before the choice-point (i.e. planning ahead of the decision – Redish, 2016), mPFC theta is more similar in frequency to dHPC theta. The finding that Re suppression did not reduce mPFC-dHPC theta coherence during non-VTE trials may also support this idea as rats were likely making their choice earlier in the stem. Furthermore, recent work from our lab supports this notion, as groups of mPFC neurons strongly predict a future choice when rats are proficiently performing a spatial working memory task (Stout and Griffin, 2020). Thus, we propose that Re suppression did not affect theta synchrony during non-VTE choice-trials as rats made their decision earlier in the stem (even if that choice was incorrect). However, when rats attempted to ‘connect’ the executive functioning (mPFC) and spatial-memory systems (HPC) under Re suppression, mPFC theta was slower in frequency, and deliberations were often associated with incorrect choices.

It should be noted that while these data suggest that Re inactivation reduces successful deliberation, it is possible that Re inactivation specifically impaired the encoding of each trajectory (Maisson et al., 2018). Failed encoding of trial-specific information would result in failed deliberation, and still fit with our synchrony results. Nonetheless, these findings lead us to wonder if a competing brain system dominates the decision-process when the Re is inactivated. For example, Packard et al., 1989 found that fornix lesions facilitated the learning of a striatal dependent task, suggesting that competition exists between the hippocampus and striatum in the intact brain. Therefore, it is possible that by preventing spatial-memory access to the mPFC (disconnecting the Re and therefore communication between the mPFC and dHPC), the competing striatal system took over, causing habit-like decision-making (i.e. perseverations). These data are in direct support of this idea as we found that perseverative errors occurred during early learning of the spatial alternation rule, disappeared with acquisition, and re-appeared in greater magnitude under Re suppression. Future studies could compare the impact of the Re on deliberation during striatal- and hippocampal-dependent tasks.

In conclusion, we examined the impact of Re suppression on VTE behaviors to answer the long-standing question of whether prefrontal-hippocampal communication supports deliberation in the rat. By inactivating the Re, a ventral midline thalamic region that connects the mPFC and HPC, we show that rats attempted to deliberate, but deliberation did not help them make a correct choice. These failed deliberations coincided with reduced prefrontal-hippocampal theta coherence, which is explained by the removal of a strong influence of the Re on the prefrontal theta oscillation. In conclusion, this study provides crucial insight to the contribution of the prefrontal-thalamo-hippocampal circuit to deliberation during spatial working memory.

## Acknowledgements

We would like to acknowledge David Redish for his contribution to VTE behavior identification and extraction as well as comments on an earlier version of this manuscript. Additionally, we would like to thank Jesse Miles and the rest of Sheri Mizumori’s lab for insightful communication about VTE behaviors in the context of our labs’ shared interests. This work could not have been made possible without the University of Delaware animal care and veterinary staff. We thank E. Myhre, B. Emanuel, D. Layfield, M. Donahue, M. Patel, L. Moedinger, A. Hallock, G. Watson, V. Chandrasekhar, and S. Amos for behavioral and technical assistance. Finally, the rat brain images used in **Fig. 1** was downloaded from SciDraw.io and was created by Gil Costa and Federico Claudi, respectively.

## Author Contributions

J.J.S and A.L.G developed the central question. J.J.S and S.A.A conceptualized analysis and questions, performed analyses and generated figures. H.L.H collected the data and provided critical conceptual contributions to the conclusions and analyses. J.J.S, H.L.H, S.A.A, and A.L.G wrote the manuscript.

## References

Allen, T. A., Narayanan, N. S., Kholodar-Smith, D. B., Zhao, Y., Laubach, M., & Brown, T. H. (2008). Imaging the spread of reversible brain inactivations using fluorescent muscimol. Journal of neuroscience methods, 171(1), 30–38.

Amemiya, S., & Redish, A. D. (2018). Hippocampal theta-gamma coupling reflects state-dependent information processing in decision making. Cell reports, 22(12), 3328–3338.

Amsel, A. (1993). Hippocampal function in the rat: cognitive mapping or vicarious trial and error?. Hippocampus, 3(3), 251–256.

Benchenane, K., Peyrache, A., Khamassi, M., Tierney, P. L., Gioanni, Y., Battaglia, F. P., & Wiener, S. I. (2010). Coherent theta oscillations and reorganization of spike timing in the hippocampal-prefrontal network upon learning. Neuron, 66(6), 921–936.

Bender, F., Gorbati, M., Cadavieco, M. C., Denisova, N., Gao, X., Holman, C., … & Ponomarenko, A. (2015). Theta oscillations regulate the speed of locomotion via a hippocampus to lateral septum pathway. Nature communications, 6(1), 1–11.

Bett, D., Allison, E., Murdoch, L. H., Kaefer, K., Wood, E. R., & Dudchenko, P. A. (2012). The neural substrates of deliberative decision making: contrasting effects of hippocampus lesions on performance and vicarious trial-and-error behavior in a spatial memory task and a visual discrimination task. Frontiers in behavioral neuroscience, 6, 70.

Bimonte, H. A., & Denenberg, V. H. (2000). Sex differences in vicarious trial-and-error behavior during radial arm maze learning. Physiology & behavior, 68(4), 495–499.

Bygrave, A. M., Jahans-Price, T., Wolff, A. R., Sprengel, R., Kullmann, D. M., Bannerman, D. M., & Kätzel, D. (2019). Hippocampal–prefrontal coherence mediates working memory and selective attention at distinct frequency bands and provides a causal link between schizophrenia and its risk gene GRIA1. Translational psychiatry, 9(1), 1–16.

Churchwell, J. C., & Kesner, R. P. (2011). Hippocampal-prefrontal dynamics in spatial working memory: interactions and independent parallel processing. Behavioural brain research, 225(2), 389–395.

Dolleman-van der Weel, M. J., Griffin, A. L., Ito, H. T., Shapiro, M. L., Witter, M. P., Vertes, R. P., & Allen, T. A. (2019). The nucleus reuniens of the thalamus sits at the nexus of a hippocampus and medial prefrontal cortex circuit enabling memory and behavior. Learning & Memory, 26(7), 191–205.

Eichenbaum, H. (2017). Prefrontal–hippocampal interactions in episodic memory. Nature Reviews Neuroscience, 18(9), 547–558.

Ferraris, M., Ghestem, A., Vicente, A. F., Nallet-Khosrofian, L., Bernard, C., & Quilichini, P. P. (2018). The nucleus reuniens controls long-range hippocampo–prefrontal gamma synchronization during slow oscillations. Journal of Neuroscience, 38(12), 3026–3038.

Gardner, R. S., Uttaro, M. R., Fleming, S. E., Suarez, D. F., Ascoli, G. A., & Dumas, T. C. (2013). A secondary working memory challenge preserves primary place strategies despite overtraining. Learning & Memory, 20(11), 648–656.

Griffin, A. L. (2015). Role of the thalamic nucleus reuniens in mediating interactions between the hippocampus and medial prefrontal cortex during spatial working memory. Frontiers in systems neuroscience, 9, 29.

Hallock, H. L., Wang, A., Shaw, C. L., & Griffin, A. L. (2013). Transient inactivation of the thalamic nucleus reuniens and rhomboid nucleus produces deficits of a working-memory dependent tactile-visual conditional discrimination task. Behavioral neuroscience, 127(6), 860.

Hallock, H. L., Wang, A., & Griffin, A. L. (2016). Ventral midline thalamus is critical for hippocampal–prefrontal synchrony and spatial working memory. Journal of Neuroscience, 36(32), 8372–8389.

Hasz, B. M., & Redish, A. D. (2020). Spatial encoding in dorsomedial prefrontal cortex and hippocampus is related during deliberation. Hippocampus, 30(11), 1194–1208.

Hembrook, J. R., & Mair, R. G. (2011). Lesions of reuniens and rhomboid thalamic nuclei impair radial maze win shift performance. Hippocampus, 21(8), 815–826.

Hembrook, J. R., Onos, K. D., & Mair, R. G. (2012). Inactivation of ventral midline thalamus produces selective spatial delayed conditional discrimination impairment in the rat. Hippocampus, 22(4), 853–860.

Hinman, J. R., Penley, S. C., Long, L. L., EscabÍ, M. A., & Chrobak, J. J. (2011). Septotemporal variation in dynamics of theta: speed and habituation. Journal of neurophysiology, 105(6), 2675–2686.

Hu, D. A. N., & Amsel, A. (1995). A simple test of the vicarious trial-and-error hypothesis of hippocampal function. Proceedings of the National Academy of Sciences, 92(12), 5506–5509.

Hyman, J. M., Zilli, E. A., Paley, A. M., & Hasselmo, M. E. (2010). Working memory performance correlates with prefrontal-hippocampal theta interactions but not with prefrontal neuron firing rates. Frontiers in integrative neuroscience, 4, 2.

Ito, H. T., Zhang, S. J., Witter, M. P., Moser, E. I., & Moser, M. B. (2015). A prefrontal– thalamo–hippocampal circuit for goal-directed spatial navigation. Nature, 522(7554), 50–55.

Johnson, Adam, and A. David Redish. “Neural ensembles in CA3 transiently encode paths forward of the animal at a decision point.” Journal of Neuroscience 27.45 (2007): 12176–12189.

Janabi-Sharifi, F., Hayward, V., & Chen, C. S. (2000). Discrete-time adaptive windowing for velocity estimation. IEEE Transactions on control systems technology, 8(6), 1003–1009.

Jones, M. W., & Wilson, M. A. (2005). Theta rhythms coordinate hippocampal–prefrontal interactions in a spatial memory task. PLoS biol, 3(12), e402.

Kastner, D. B., Gillespie, A. K., Dayan, P., & Frank, L. M. (2020). Memory Alone Does Not Account for the Way Rats Learn a Simple Spatial Alternation Task. Journal of Neuroscience, 40(38), 7311–7317.

Kidder, K. S., Miles, J. T., Baker, P. M., Hones, V. I., Gire, D. H., & Mizumori, S. J. (2021). A selective role for the mPFC during choice and deliberation, but not spatial memory retention over short delays. Hippocampus.

Kropff, E., Carmichael, J. E., Moser, E. I., & Moser, M. B. (2021). Frequency of theta rhythm is controlled by acceleration, but not speed, in running rats. Neuron, 109(6), 1029–1039.

Layfield, D. M., Patel, M., Hallock, H., & Griffin, A. L. (2015). Inactivation of the nucleus reuniens/rhomboid causes a delay-dependent impairment of spatial working memory. Neurobiology of learning and memory, 125, 163–167.

Lee, I., & Kesner, R. P. (2003). Time-dependent relationship between the dorsal hippocampus and the prefrontal cortex in spatial memory. Journal of Neuroscience, 23(4), 1517–1523.

Maisson, D. J. N., Gemzik, Z. M., & Griffin, A. L. (2018). Optogenetic suppression of the nucleus reuniens selectively impairs encoding during spatial working memory. Neurobiology of Learning and Memory, 155, 78–85.

McFarland, W. L., Teitelbaum, H., & Hedges, E. K. (1975). Relationship between hippocampal theta activity and running speed in the rat. Journal of comparative and physiological psychology, 88(1), 324.

eyer-Mueller, C., Jacob, P. Y., Montenay, J. Y., Poitreau, J., Poucet, B., & Chaillan, F. A. (2020). Dorsal, but not ventral, hippocampal inactivation alters deliberation in rats. Behavioural brain research, 390, 112622.

Mitra, P. (2007). Observed brain dynamics. Oxford University Press.

uenzinger, K. F., & Gentry, E. (1931). Tone discrimination in white rats. Journal of Comparative Psychology, 12(2), 195.

Muenzinger, K. F. (1938). Vicarious trial and error at a point of choice: I. A general survey of its relation to learning efficiency. The Pedagogical Seminary and Journal of Genetic Psychology, 53(1), 75–86.

O‘Neill, P. K., Gordon, J. A., & Sigurdsson, T. (2013). Theta oscillations in the medial prefrontal cortex are modulated by spatial working memory and synchronize with the hippocampus through its ventral subregion. Journal of Neuroscience, 33(35), 14211–14224.

Packard, M. G., Hirsh, R., & White, N. M. (1989). Differential effects of fornix and caudate nucleus lesions on two radial maze tasks: evidence for multiple memory systems. Journal of Neuroscience, 9(5), 1465–1472.

Papale, A. E., Stott, J. J., Powell, N. J., Regier, P. S., & Redish, A. D. (2012). Interactions between deliberation and delay-discounting in rats. Cognitive, Affective, & Behavioral Neuroscience, 12(3), 513–526.

Paxinos, G., & Watson, C. (2006). The rat brain in stereotaxic coordinates: hard cover edition. Elsevier.

Redish, A. D. (2016). Vicarious trial and error. Nature Reviews Neuroscience, 17(3), 147.

Santos-Pata, D., & Verschure, P. F. (2018). Human vicarious trial and error is predictive of spatial navigation performance. Frontiers in behavioral neuroscience, 12, 237.

Schmidt, B., Duin, A. A., & Redish, A. D. (2019). Disrupting the medial prefrontal cortex alters hippocampal sequences during deliberative decision making. Journal of neurophysiology, 121(6), 1981–2000.

Schmidt, B., Papale, A., Redish, A. D., & Markus, E. J. (2013). Conflict between place and response navigation strategies: effects on vicarious trial and error (VTE) behaviors. Learning & Memory, 20(3), 130–138.

Sigurdsson, T., Stark, K. L., Karayiorgou, M., Gogos, J. A., & Gordon, J. A. (2010). Impaired hippocampal–prefrontal synchrony in a genetic mouse model of schizophrenia. Nature, 464(7289), 763–767.

Sławińska, U., & Kasicki, S. (1998). The frequency of rat’s hippocampal theta rhythm is related to the speed of locomotion. Brain research, 796(1-2), 327–331.

Tolman, E. C. (1939). Prediction of vicarious trial and error by means of the schematic sowbug. Psychological Review, 46(4), 318.

Tolman, E. C. (1948). Cognitive maps in rats and men. Psychological review, 55(4), 189.

arela, C., Kumar, S., Yang, J. Y., & Wilson, M. A. (2014). Anatomical substrates for direct interactions between hippocampus, medial prefrontal cortex, and the thalamic nucleus reuniens. Brain Structure and Function, 219(3), 911–929.

Vertes, R. P., Hoover, W. B., Do Valle, A. C., Sherman, A., & Rodriguez, J. J. (2006). Efferent projections of reuniens and rhomboid nuclei of the thalamus in the rat. Journal of comparative neurology, 499(5), 768–796.

Vertes, R. P., Hoover, W. B., Szigeti-Buck, K., & Leranth, C. (2007). Nucleus reuniens of the midline thalamus: link between the medial prefrontal cortex and the hippocampus. Brain research bulletin, 71(6), 601–609.

Viena, T. D., Linley, S. B., & Vertes, R. P. (2018). Inactivation of nucleus reuniens impairs spatial working memory and behavioral flexibility in the rat. Hippocampus, 28(4), 297–311.

Voss, J. L., Warren, D. E., Gonsalves, B. D., Federmeier, K. D., Tranel, D., & Cohen, N. J. (2011). Spontaneous revisitation during visual exploration as a link among strategic behavior, learning, and the hippocampus. Proceedings of the National Academy of Sciences, 108(31), E402–E409.

Voss, J. L., & Cohen, N. J. (2017). Hippocampal cortical contributions to strategic exploration during perceptual discrimination. Hippocampus, 27(6), 642–652.

Wang, G. W., & Cai, J. X. (2006). Disconnection of the hippocampal–prefrontal cortical circuits impairs spatial working memory performance in rats. Behavioural brain research, 175(2), 329–336.

Wang, G. W., & Cai, J. X. (2008). Reversible disconnection of the hippocampal-prelimbic cortical circuit impairs spatial learning but not passive avoidance learning in rats. Neurobiology of learning and memory, 90(2), 365–373.

